# 3D Multiplexed Immunohistochemistry Using Opal-TSA Amplification for Enhanced Imaging of Pulmonary TB Lesions

**DOI:** 10.1101/2024.10.23.619885

**Authors:** Suruchi Lata, Shivraj M. Yabaji, Aoife K O’Connell, Hans P Gertje, Nicholas A Crossland, Igor Kramnik

## Abstract

*Mycobacterium tuberculosis* hijacks the host immune system and persists for several years before the onset of active disease. Spatial characterization of epithelial compartments, immune cell populations and bacteria simultaneously within tissue specimens provide significant information about host pathogen interactions. Here, we present a protocol to detect multiple protein markers using Opal™-TSA conjugated fluorescent dyes in free floating 10% neutral buffered formalin fixed thick tissue sections (50-100 μm), with minimal additional tissue processing not requiring specialized equipment. Use of thick sections provides more information as compared to classic thin microtomy sections (3-10 μm), including the capacity for Z stacking and three-dimensional image rendering. Importantly, reduced tissue processing of samples with this method preserves endogenous fluorescent reporter signal. Use of Opal™-TSA conjugated fluorescent dyes enhances the sensitivity of low expressing proteins and supports the use of primary antibodies raised in the same species.

**Before you begin:** This protocol describes the specific use of Mtb infected mouse lungs, but is applicable to any tissue type and species of origin.

**Institutional permissions:** Obtain institutional permission to perform animal studies and collect tissues under an approved Institutional Animal Care and Use Committee (IACUC) or Institutional Review Board protocol. Our protocol was approved by Boston University’s Institutional Animal Care and Use Committee (IACUC protocol number PROTO201800218).

**Mice:** B6J.C3-*Sst1*^*C3HeB/Fej*^ Krmn and B6. Sst1S, ifnb-YFP mice were developed in our laboratory (available from MMRRC stock # 043908-UNC).

**Highlights:**

- Compatible with Opal™-TSA conjugated fluorescent dyes resulting in enhanced signal sensitivity with low background noise
- Reduced tissue processing preserves endogenous fluorescent reporter signals
- Compatible with primary antibodies raised in the same species
- Doesn’t require specialized automated instruments

**Graphical Abstract:** 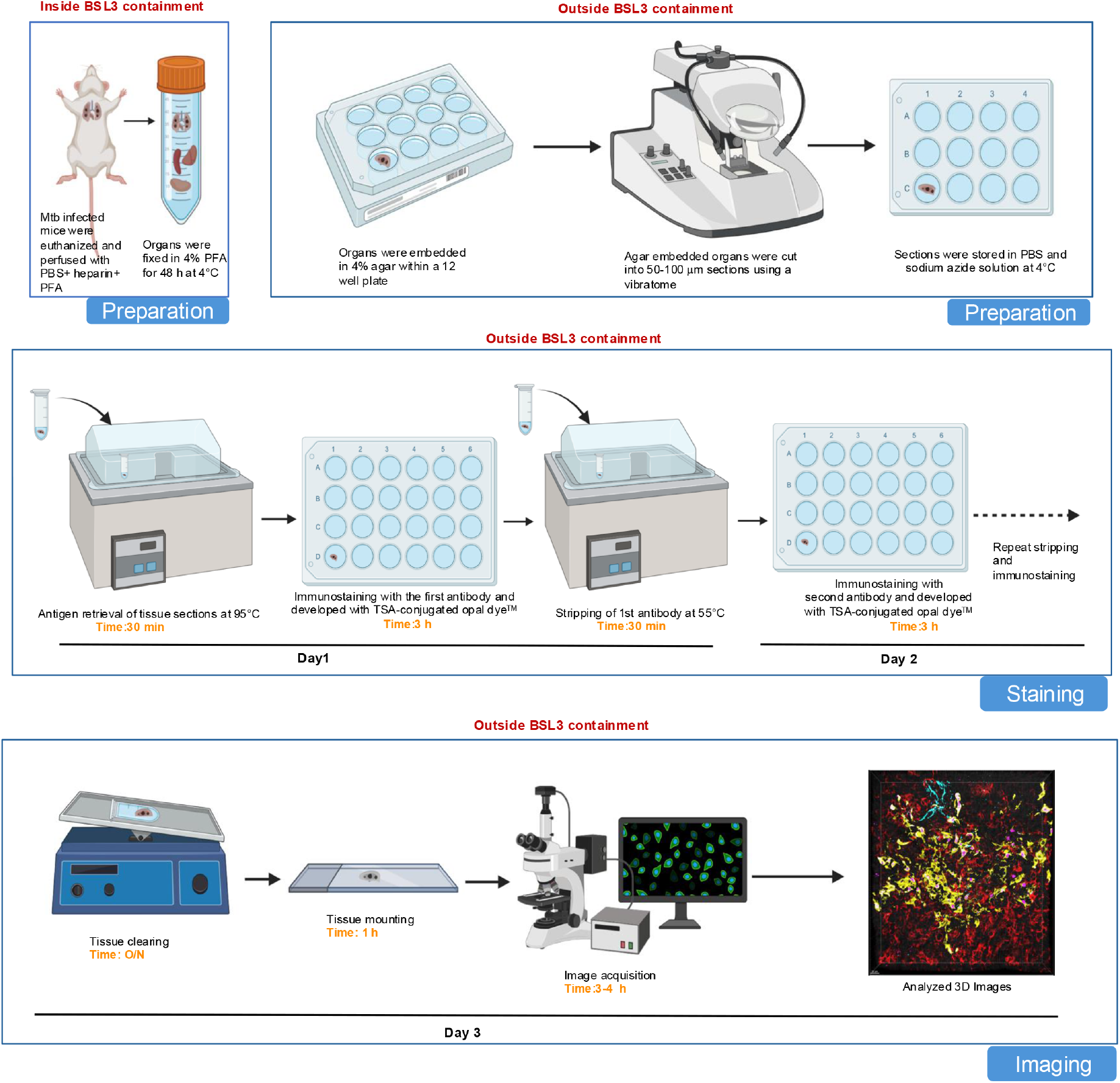

## Background

*Mycobacterium tuberculosis* (Mtb) resides inside the host for several years before the onset of active disease. Mtb is primarily deposited to the lung through aerosolized droplets and phagocytized by alveolar macrophages. Meanwhile the host’s immune cells migrate to the site of infection and try to clear the infection. Mtb resides inside the lung by forming discrete encapsulated granulomas, which is the hallmark of tuberculosis. The granuloma is composed of both myeloid and lymphoid cells surrounded by a fibrotic capsule. Mtb survives within the granuloma by escaping host immune surveillance and coping with stresses like hypoxia, low pH and nutrient sequestration. To study the host pathogen interactions during tuberculosis, the spatial distribution of immune cells and Mtb within granulomas at different stages of disease progression could provide a solid framework for temporal granuloma niche evolution. In this study we have used B6. Sst1S mouse, which develops human like pulmonary TB including formation of caseating granulomas (Pichugin et al., 2009). We used thick tissue sections (50-100 μm) derived from infected mouse lungs for immunostaining with 3-4 protein markers simultaneously. Use of multiple markers at one time in thick sections provides more information, while preserving precious resources. Use of Opal™-TSA conjugated fluorescent dyes is compatible with utilization of primary antibodies raised in the same species of origin (Stack et al., 2014; Toth and Mezey, 2007). Although Opal dyes™ have been routinely used for fluorescent multiplex IHC (fmIHC) on FFPE tissue sections. Here we utilize Opal dyes™ on 10% neutral buffered formalin fixed thick tissue sections and found that iterative stripping and deposition of these dyes is compatible which thick tissue sections, results in enhanced signal sensitivity, with less background even at low magnification, which is also helpful in identifying regions of interest for subsequent high magnification imaging.

### Overview of TSA mediated immunostaining method

We use florescent labeled Opal dyes™ for antigen detection, which utilizes tyramide signal amplification (TSA) to amplify the signal detection. Horseradish peroxidase (HRP) conjugated to the secondary antibody catalyzes the covalent deposition of the fluorophore to tyrosine residues in tissues in close proximity of the target antigen in the presence of a low concentration of hydrogen peroxide (**Fig.1**).

**Figure 1:**
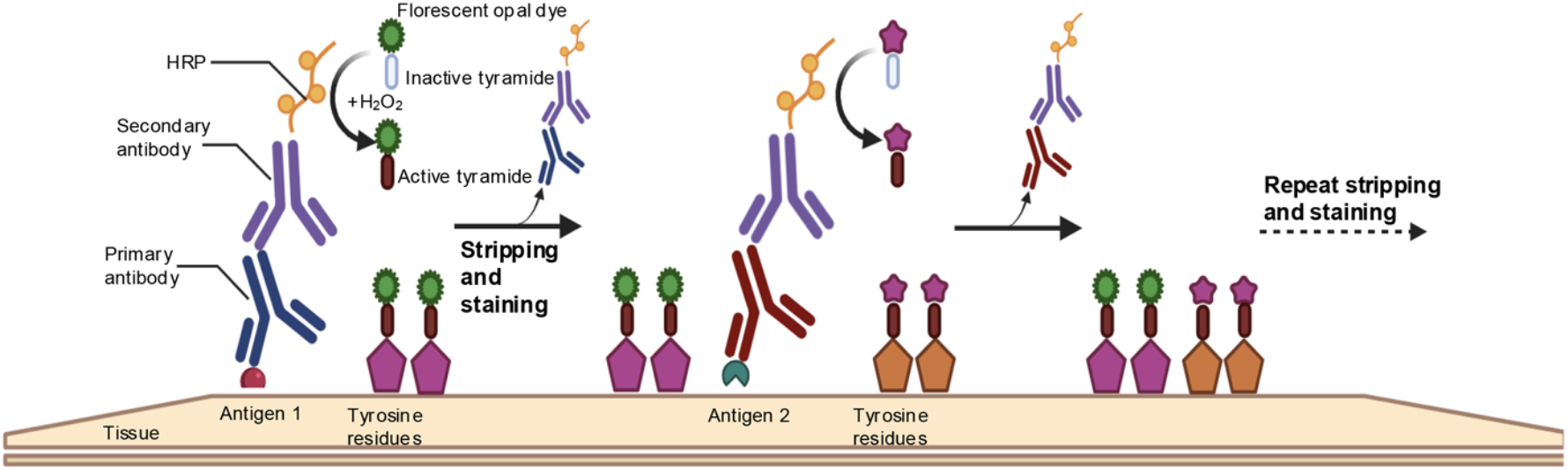
Mechanism of tyramide signal amplification (TSA) staining. Primary antibodies recognize the epitope on the target protein. HRP conjugated secondary antibodies recognize the species-specific Fc region of the primary antibodies. HRP catalyzes the conversion of fluorescently labeled tyramide to active tyramide in the presence of hydrogen peroxide, which covalently binds to tyrosine residues in spatial proximity to the target protein.

### Sample preparation

#### Anesthesia and perfusion

**Timing:** 2 days

This protocol is optimized for Mtb infected mice. Mice are anesthetized by injecting ketamine-xylazine solution intraperitoneally. Perfusion was conducted with a perfusion pump containing 1X PBS with 10 IU/mL heparin (in one tube and 10% neutral buffered formalin in another tube. Lungs were inflated to normal physiologic volume before harvesting and kept in 10% formalin solution for 48 h at 4°C before removing from BSL3. Organs were transferred to PBS and stored at 4°C.

#### Key resources table

| REAGENT or RESOURCE | SOURCE | IDENTIFIER |
| --- | --- | --- |
| **Antibodies** |  |  |
| Rabbit anti-P63 (1:100) | Cell Signaling Technology | Cat# 13109T<br>RRID:AB_2637091 |
| Rabbit anti-Thyroid Transcription factor-1 (TTF-1) 1:100 | Abcam | Cat# ab76013,<br>RRID:AB_1310784 |
| Rabbit anti- iNOS (1:100) | Abcam | Cat# ab15323,<br>RRID:AB_301857 |
| Rabbit anti-Ki67 (1:100) | Cell Signaling Technology | Cat# 12202,<br>RRID:AB_2620142 |
| Rabbit anti-ProSPC (1:250) | Seven Hills Bioreagents | Cat# WRAB-9337,<br>RRID:AB_2335890 |
| Rabbit anti-Keratin 17 (1:50) | Cell Signaling Technology | Cat# 4543,<br>RRID:AB_2133014 |
| Rabbit anti-Keratin 8 (1:50) | Invitrogen | Cat# MA5-32118,<br>RRID:AB_2809410 |
| Rabbit anti-Smooth Muscle Action (1:200) | Cell Signaling Technology | Cat# 19245,<br>RRID:AB_2734735 |
| Rabbit anti-GFP (1:100) | Invitrogen | Cat#A11122,<br>RRID:AB_221569 |
| **Bacterial strain** |  |  |
| *Mycobacterium tuberculosis* Erdman strain (SSB-GFP, *smyc'*::mCherry) | (Giacalone et al., 2021; Sukumar et al., 2014) | N/A |
| **Biological samples** |  |  |
| Mouse: B6J.C3-Sst1<sup>C3HeB/Fej</sup>Krmn | (Pichugin et al., 2009) | Stock No: 043908-UNC<br>https://www.mmrrc.org |
| Mouse: B6. Sst1S, ifnb-YFP | (Scheu et al., 2008; Yabaji et al., 2023) | N/A |
| **Chemicals, peptides, and recombinant proteins** |  |  |
| Gibco™ DPBS | Fisher scientific | Cat#14-190-235 |
| Heparin | Thermo Scientific | Cat# A16198.03 |
| Paraformaldehyde | Thermo Scientific | Cat# J61899.AP |
| Agar | Fisher scientific | Cat# BP1423-500 |
| Sodium azide | Sigma Aldrich | Cat# S2002-25G |
| Tris based buffer-Cell Conditioning 1 (CC1) | Roche | Cat# 950-124 |
| Hydrogen peroxide | Fisher Scientific | Cat# H325500 |
| 1X Antibody Diluent/ Block | Akoya Biosciences | Cat#ARD1001EA |
| Ventana Antibody Diluent with Casein | Roche | Cat# 06440002001 |
| ImmPRESS HRP Anti-Rabbit IgG (Peroxidase) Polymer Detection Kit, made in Goat | Vector Laboratories | Cat#MP-7451, |
| Opal™ 540 Reagent Pack | Akoya Biosciences | FP1494001KT |
| Opal™ 570 Reagent Pack | Akoya Biosciences | FP1488001KT |
| Opal™ 620 Reagent Pack | Akoya Biosciences | FP1495001KT |
| Opal™ 650 Reagent Pack | Akoya Biosciences | FP1496001KT |
| 1X Plus Automation Amplification Diluent | Akoya Biosciences | FP1609 |
| 2- Mercaptoethanol | MP Biomedicals | 190242 |
| Sodium Dodecyl sulfate | Fisher Scientific | BP2436200 |
| 0.5 M Tris-HCL (pH 6.8) | Fisher Scientific | J12112-36 |
| Histodenz | Sigma Aldrich | Cat# NC1589271 |
| N-METHYLACETAMIDE | Sigma Aldrich | Cat# 50-183-6896 |
| Triton X-100 | Sigma Aldrich | Cat# T8787 |
| 1-Thioglycerol | Sigma Aldrich | Cat# 50-184-5430 |
| Vectashield antifade mounting media | Vector Laboratories | Cat# NC1695563 |
| **Software** |  |  |
| Imaris Viewer | Oxford Instruments | [https://imaris.oxinst.com/microscopy-imaging-software-free-trial?](https://imaris.oxinst.com/microscopy-imaging-software-free-trial?source=viewer)<br>source=viewer |
| **Other** |  |  |
| LEICA VT1200 S fully automatic vibratome | Leica Biosystems | N/A |
| double edge stainless steel blades | Personna | N/A |
| Fisher brand super frost microscope slides (75 X 25 mm) | Fisher Scientific | 12-550-15 |
| Microscope cover glass, No.1 (24 X 50 mm) | Globe scientific INC | 1415-10 |
| Leica SP5 confocal microscope | Leica Biosystems | N/A |
| Water bath | Fisher Scientific | 15-462-2 |
| Rocking shaker | Fisher Scientific | 260100F |
| Clear nail polish | Ted Pella | N/A |

## Materials and equipment

### Heparin solution

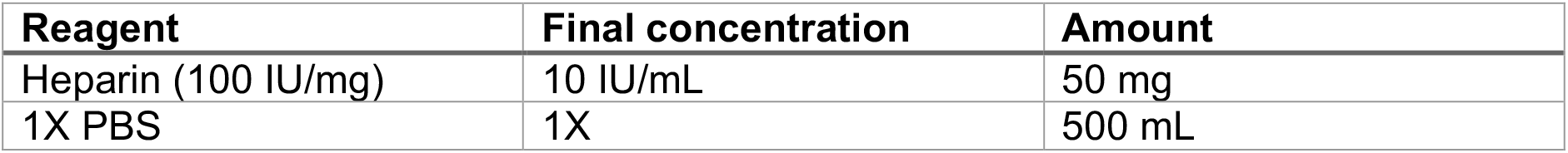

Materials and equipment Heparin solution
| Reagent | Final concentration | Amount |
| --- | --- | --- |
| Heparin (100 IU/mg) | 10 IU/mL | 50 mg |
| 1X PBS | 1X | 500 mL |

#### Stripping buffer

| Reagent | Final concentration | Amount |
| --- | --- | --- |
| BME | 0.8% | 0.8 mL |
| SDS | 10% | 20 mL |
| Tris-HCL (pH 6.8) | 0.5M | 12.5 mL |

Adjust volume to 100 mL with deionized water.

**Critical**-Prepare under a fume hood

#### Ce3D clearing solution

| Reagent | Amount (5 mL) |
| --- | --- |
| 40% N-METHYLACETAMIDE | 2 mL |
| Histodenz | 4 g |
| 40% N-METHYLACETAMIDE | 750 μL |
| Triton X-100 | 5 μL |
| 1-Thioglycerol | 25 μL |

**Step-by-step method details**

#### Outline of the protocol

Step-by-step method details Outline of the protocol
| Step |  | Time | Temperature | Containment |
| --- | --- | --- | --- | --- |
| 1. | Embedding of organ | 2 h | 4°C | BSL2 |
| 2. | Tissue sectioning | 1 h | RT | BSL2 |
| 3. | Washing x 3 | 30 min | RT | BSL2 |
| 4. | Antigen retrieval and permeabilization | 30 min | 95°C | BSL2 |
| 5. | Washing x 3 | 30 min | RT | BSL2 |
| 6. | Blocking | 12 min | RT | BSL2 |
| 7. | Primary antibody | 1 h | 37°C | BSL2 |
| 8. | Washing x 3 | 30 min | 37°C | BSL2 |
| 9. | Blocker | 8 min | RT | BSL2 |
| 10. | HRP conjugated secondary antibody | 20 min | 37°C | BSL2 |
| 11. | Washing x 3 | 30 min | 37°C | BSL2 |
| 12. | Opal™ fluorophore | 20 min | RT | BSL2 |
| 13. | Stripping | 30 min | 56°C | BSL2 |
| 14. | Washing | 40 min | RT | BSL2 |
|  | **Repeat staining with second antibody** |  |  |  |
| 15. | Nuclear staining | 10 min | RT | BSL2 |
| 16. | Tissue clearing | 10-12 h | RT | BSL2 |
| 17. | Mounting of sections | 1-2 h | RT | BSL2 |
| 18. | Image acquisition | 4-5 h | RT | BSL2 |

Mount tissues on microscopic slides

### Day 1: Preparation of tissue sections

**Timing:** 1 h+

This protocol is optimized with Mtb infected mouse lungs but can be used with other organs and species as well.

1. To embed a fixed lung lobe, pour around 1mL of 4% melted agar in PBS into a 12 well plate and put the lung lobe on the top, pour around 1mL of melted agar on top of the lung again to embed the lung completely. Use forceps to reposition the lung lobe in the agar if needed. Allow the sample to solidify completely at 4°C for around 2 h in the dark to preserve endogenous fluorescent signals before starting the sectioning process.
2. Trim the extra agar from the edges of the block and secure it to the specimen disc of the vibratome with super glue.
3. Cut 50-100 μm sections (0.145 mm/s, 70 Hz, blade angle 5°), and place them into cold PBS(Giacalone et al., 2021; Tan et al., 2013). Tissue sections can be stored in PBS with 0.01% sodium azide for up to 3-5 months at 4°C in the dark

### Day 1: Washing of the tissue sections

**Timing: 30 min**

4. Wash tissue section in a non-adherent petri plate containing PBS solution on a nutator mixer for 5 min at RT. Repeat the washing step in fresh PBS for 2 times. Hold the tissue sections carefully with a disposable inoculating loop or paint brush while transferring.

### Day 1: Antigen retrieval

**Timing: 45 min**

Protein crosslinking during fixation process masks the epitopes and may reduce antigen-antibody binding efficiency. The antigen retrieval process unmasks the epitopes and enables the antibodies to recognize the target epitopes within the tissue specimen. This process can be enzymatic (PIER), or heat induced (HIER). Here we used the heat-induced epitope retrieval method (Shi et al., 1991).

**Alternatives**: Permeabilize the tissue sections by incubating them in 2% triton X 100 for 1 h for similar results without compromising tissue integrity.

5. Transfer 5-6 tissue sections into a microfuge containing 300 μL of Tris/Borate/EDTA buffer solution (CC1; pH 8, Roche) and incubate for 30 min at 95°C in a water bath. The volume of the buffer solution should be sufficient to submerge the tissue sections.
6. Keep the sample at room temperature to cool down completely and then transfer the tissues carefully to the 24 well plate. Depending on the size of the tissue place 3-5 tissues in each well.
7. Wash the tissue sections for 5 min with 200 μL of reaction buffer (Roche) at room temperature on a nutator shaker. Repeat the washing process twice. Transfer the tissue carefully to a new well after every wash with forceps

### Day 1: Blocking of endogenous peroxidase activity

**Timing: 22 min**

Endogenous peroxidase activity can react with hydrogen peroxide and catalyze tyramide signal amplification (TSA) which leads to nonspecific staining. Hence it is important to block endogenous peroxidase activity before starting the staining process. Here we use 3% hydrogen peroxide (v/v) for blocking.

8. Submerge tissue sections in 3% hydrogen peroxide (v/v) for 12 min at room temperature under rocking condition.
9. Wash the samples with 200 μL of reaction buffer (Roche) at room temperature on a nutator shaker for 5 min. Repeat the washing process twice. Transfer the tissue carefully to a new well after washing.

### Day 1: Blocking non-specific sites

**Timing: 8 min**

Antibodies may bind to nonspecific targets. Blocking solutions minimize nonspecific binding of antibodies and reduce the background noise during fluorescent staining. Here we use 1X antibody diluent/block solution (Akoya biosciences).

10. Pour a few drops of blocking buffer to submerge the tissue sections and place the plate on a nutator shaker for 8 min at room temperature.

### Day 1: Primary antibody incubation

**Timing: 1 h**

11. Prepare the primary antibody dilution using Ventana antibody diluent with casein. Use Signal Stain^®^ Antibody Diluent (CST) for primary antibodies sourced from cell signaling technology.
11. Pipet 200 μL of diluted primary antibody into each well of a 24 well plate containing tissues. Transfer tissue sections from the blocking buffer to the primary antibody solution.
12. Place the plate on a nutator shaker set at 37°C for 1 h.
13. Wash the tissue samples for 5 min with 200 μL of reaction buffer (Roche) at room temperature on a nutator shaker. Repeat the washing process twice. Transfer the tissue carefully to a new well after every wash.

### Day 1: Blocking non-specific sites

**Timing: 8 min**

14. Pour a few drops of blocking buffer (Akoya biosciences) to submerge the tissue sections and place the plate on a nutator shaker for 8 min at room temperature.

### Day 1: Secondary antibody incubation

**Timing: 25 min**

Pipet 200 μL of secondary antibody in each well of 24 well plate. Use secondary antibody directed against the species used for the primary antibody. Here we use ImmPRESS® HRP goat anti-rabbit secondary antibody (Vector Laboratories).

15. Transfer tissue sections from the blocking buffer to the secondary antibody solution and place the plate on a rocker set at 37°C for 20 min.
16. Wash the tissue samples for 5 min with 200 μL of reaction buffer (Roche) at room temperature on a nutator shaker.

### Day 1: Opal™ fluorophore incubation

**Timing: 35 min**

Tyramide conjugated Opal™ fluorophores are used to develop primary-secondary antibody complexes. HRP molecules conjugated to the secondary antibody catalyze tyramide signal amplification (TSA) and the fluorophore is deposited near the target antigen inside the tissue via tyrosine residues. Opal™ reagents are reconstituted in 75

μLl of DMSO before use and diluted typically between 1:75 to 1:100 in 1X plus automation amplification buffer (Akoya Biosciences).

17. Pipet 200 μL of diluted Opal™ fluorophore into one well and transfer the tissue sections. Incubate the plate on a nutator shaker for 20 min at room temperature in the dark.
18. Wash the tissue samples for 5 min with 200 μL of reaction buffer (Roche) at room temperature on a nutator shaker. Repeat the washing process twice and transfer the tissue carefully into new well after every wash.

### Day 1: Antibody stripping

**Timing: 1 h**

Before starting the staining process with the second primary antibody, it is important to remove the first set of primary and secondary antibody complexes to minimize the cross reactivity and background noise. Since TSA deposits tyramide conjugated fluorophores covalently to the tyrosine side chains neighboring the target antigen, stripping of primary and secondary antibody does not remove fluorophores from the tissue specimen. Effective stripping allows for use of antibodies raised in the same species and multiplexing. IHC with FFPE tissue sections uses citric acid buffer and high temperature (100°C) for stripping of primary and secondary antibody complexes. When we conducted HIER on free floating thick tissue sections at 95°C for 30 min using citric acid buffer, we determined tissue sections are prone to damage as a consequence of exposure to iterative high temperatures. Hence, we relied on another stripping method which doesn’t require high temperature. We used the surfactant sodium dodecyl sulfate (SDS) and the reducing agent β-mercaptoethanol (BME) stripping solution to remove the first set of primary and secondary antibodies without removing the fluorophores (Ehrenberg et al., 2020).

19. Prepare BME/SDS stripping buffer within a chemical fume hood. Mix 20 mL of 10% SDS, 12.5 mL of 0.5 M Tris-HCL (pH 6.8), 0.8 mL of β-mercaptoethanol (BME) and 67.5 mL of deionized water in a chemical fume hood.
20. Pipet 1 mL of BME/SDS buffer to a 2 mL microcentrifuge and transfer the tissue sections in the tube.
21. Place the tube in a water bath set at 50°C for 30 min. Wrap the tubes with aluminum foil to protect the tissue sections from light.
22. Let the samples cool at room temperature for at least 20 min. Transfer the tissue samples to 24 well plates inside the fume hood.
23. Wash the tissue samples for 10 min with 500 μL of TBST at room temperature on a nutator. Repeat the washing process twice and transfer the tissues carefully into a new well after each wash.

**Critical**-Protect tissue sections from light. Use a chemical fume hood for stripping buffer preparation.

### Day 2: Repeat staining and stripping

**Timing:** 8-16 h

24. After the first round of primary and secondary antibody stripping, repeat the staining process with a second primary antibody. During this cycle add 1X antibody diluent/block solution (Akoya biosciences) before adding primary and secondary antibody. There is no need to block endogenous peroxidase at this point.
25. Add the HRP conjugated secondary antibody and develop the second antibody with a different Opal™-TSA conjugated fluorophore (having a different excitation and emission spectra as compared to the first fluorophore compatible with your accessible imaging modalities).
26. Repeat the staining process with all the required primary antibodies. We have used up to four primary antibodies raised within the same species using this method.

### Day 2: Nuclear staining

**Timing:** 15 min

When tissue sections are stained with all the required markers, they are incubated in Hoechst 33342 solution for nuclear detection.

27. Dilute Hoechst 33342 solution in PBS (1:1000) and pipet 500 μL in a well. Transfer tissue sections in Hoechst 33342 solution and incubate the plate at room temperature for 10 min on a nutator mixer.
28. Wash the tissue sections with 500 μL of PBS for 5 min on nutator mixer.

### Day 2: Tissue clearing(Li et al., 2019)

**Timing: Overnight**

29. Pour RapiClear®clearing solution 1.47 (SUNJin Lab) or Ce3D clearing solution (22% N-methyl acetamide, 86% Histodenz, 0.1% Triton X-100, 0.5% 1-thioglycerol) into a 2mL tube and transfer the stained tissue sections into the tube.
30. Place the tube on a nutator shaker overnight at room temperature.

**Note:** Reagents for Ce3D clearing solution must be added in the following order: 2 mL of 40% (v/v) N-methyl acetamide, 4 g Histodenz, 750 μL of 40% (v/v) N-methyl acetamide, and 5 μL of Triton X-100. Seal the tube with Parafilm. Place tube upright in a 37°C shaking incubator and shake at 150 rpm until it dissolves completely. Add 25 μL of 1-Thioglycerol and mix well on rocker. Store at room temperature. This solution can be used for up to 1 month.

**Note:** N-methyl acetamide is solid at around room temperature. To dissolve place the entire bottle of 100% N-methyl acetamide into a 37°C incubator for >1 h, until fully dissolved. Then, transfer 20 mL of 100% N-methyl acetamide to a 50 mL falcon tube with a pre-warmed 25 mL serological pipette to prevent N-methyl acetamide from sticking to the pipette wall. Add 30 mL of PBS and mix to make a 40% (v/v) stock solution.

**Critical**-N-methyl acetamide is a presumed human reproductive and fetal toxicant. Use a chemical fume hood and appropriate personal protective equipment throughout the preparation of N-methyl acetamide. Collect the waste in an appropriate container for environmental health and safety (EH&S) disposal in accordance with your institution’s regulations.

### Day 3: Mounting of tissue sections

**Timing: 30+ min**

We mount stained tissue sections on Superfrost® Plus Microscope Slides. Tissue sections adhere to the slides before mounting.

31. Cleared tissue sections are removed from the clearing solution utilizing a disposable inoculating loop and placed on the microscopic slide.
32. Carefully unfurl and reposition the tissue sections with the help of the inoculating loop. At this point if sections are too viscous to handle, add a few drops of PBS on the sections before moving the tissue with the help of inoculating loop.
33. Gently move the tissue with the inoculating loop so that it lies completely flat on the slides. Gently pat the tissues sections with Kim wipes and wipe out the excess PBS solution completely from the slide.

**Note**: The tissue sections should be unfurled as quickly as possible if PBS solution has been added because it can reverse the clearing process. In place of an inoculating loop, one can use a paint brush.

34. Allow the tissue sections to dry on the positively charged surface of the microscopic slides to enable tissue to adhere at room temperature in the dark.
35. Pipette a few drops of vectashield antifade mounting media over the tissue and gently place a coverslip (No. 1; thickness 150μm). Remove trapped air bubbles by pressing the coverslip gently with forceps.
36. Allow the cover slipped slides to dry in the dark for 30 min and apply clear nail polish around the edges of the coverslip to seal the gap between the coverslip and the slide.
37. Slides are ready for imaging and can be stored at 4°C away from light.

### Day 3: Image acquisition

Images can be acquired using epi-fluorescent microscopes with appropriate emission light sources and filters cubes or confocal microscopes with appropriate excitation and emission laser lines according to the Opal dyes™ used for immunostaining. In this study we used a Leica SP5 confocal microscope for image acquisition enalbing generation of Z-stacks. Imaris 10.1.0 was used for 3D reconstruction of confocal images.

### Expected outcomes

This protocol results in imaging of thick tissue section with 4-5 markers at a time (**Fig.2 and 3**). Use of thick sections and 3D reconstruction of images provides more information as compared to thin sections. Compatibility of this method with Opal dyes™ and the stripping process allows sequential application of primary antibodies raised from the same species. Use of Opal dyes™ results in enhanced signal sensitivity as compared to conventional immunofluorescence with minimal background noise even at low magnification (**Fig.4**), which helps to quickly detect fields of interest and provide more information. Minimum tissue processing of samples prior to sectioning preserves endogenous fluorescent reporter signals (**Fig. 5**).

**Figure 2:**
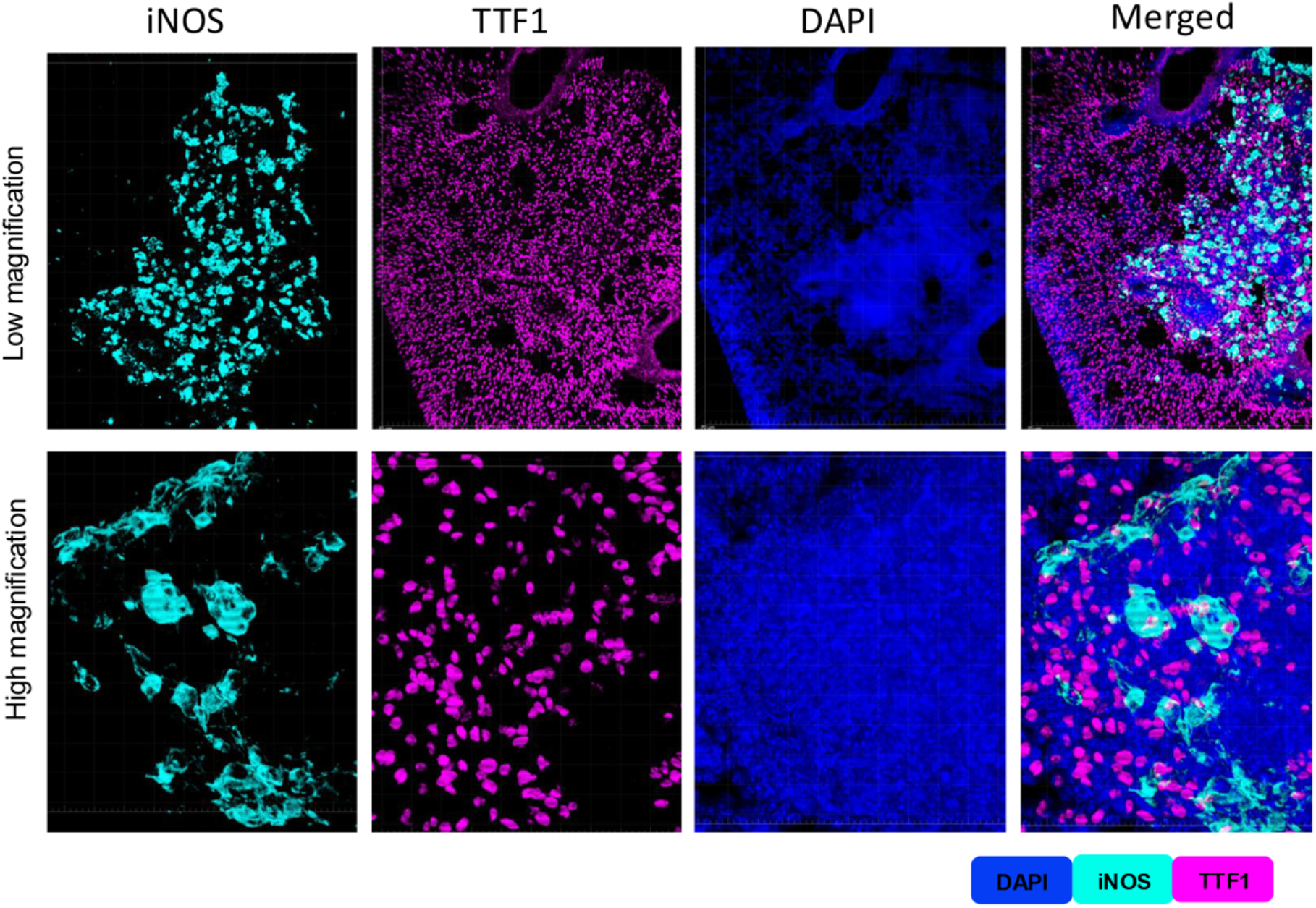
Confocal 3D images of fluorescent multiplex immunohistochemistry (fmIHC) using tyramide signaling amplification (TSA) of thick lung sections with two different primary antibodies raised in same species. Thick sections were first stained with an iNOS primary antibody (raised in rabbit) and developed with an HRP-conjugated anti-Rb secondary antibody and Opal™ 690 dye, stripped with BME/SDS stripping buffer and then immunostained with a TTF1 primary antibody (raised in rabbit) and developed with an HRP-conjugated anti-Rb secondary antibody and Opal™ 570 dye; images were acquired at low magnification (10X; upper row) and high magnification (63X; lower row). Images highlight the absence of bleed through between the two channels supporting the efficacy of our stripping method. iNOS shown in teal and TTF1 shown in magenta.

**Figure 3:**
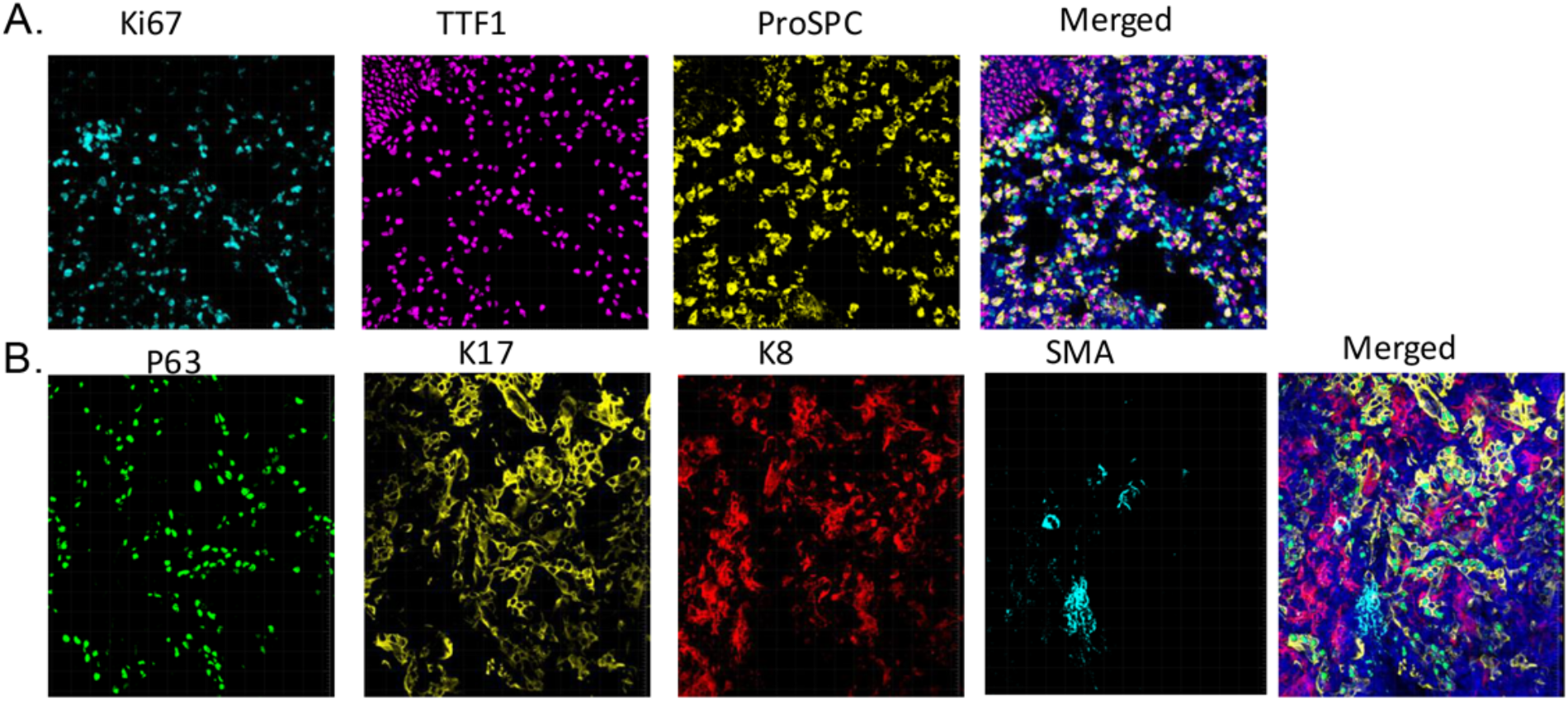
Confocal 3D images of fluorescent multiplex immunohistochemistry (fmIHC) using tyramide signaling amplification (TSA) labelling of thick lung sections with different antibodies raised in the same species. Thick sections were stained with different antibodies (all raised in rabbit) and developed with different opal dyes™. **A**. Three different antibodies were used: Ki67 was developed with opal™ 650 dye (teal), TTF1 was developed with opal™ 620 dye (magenta) and ProSPC was developed with opal™ 540 dye (yellow). **B**. four different antibodies were used: P63 was developed with opal™ 570 dye (green), K17 was developed with opal™ 540 dye (yellow), K8 was developed with opal™ 620 dye (red) and smooth muscle actin (SMA) was developed with opal™ 650 dye (teal). Images showing that there is no bleed through between the channels and stripping method successfully removes one set of antibodies. Images were acquired at 40X magnification.

**Figure 4:**
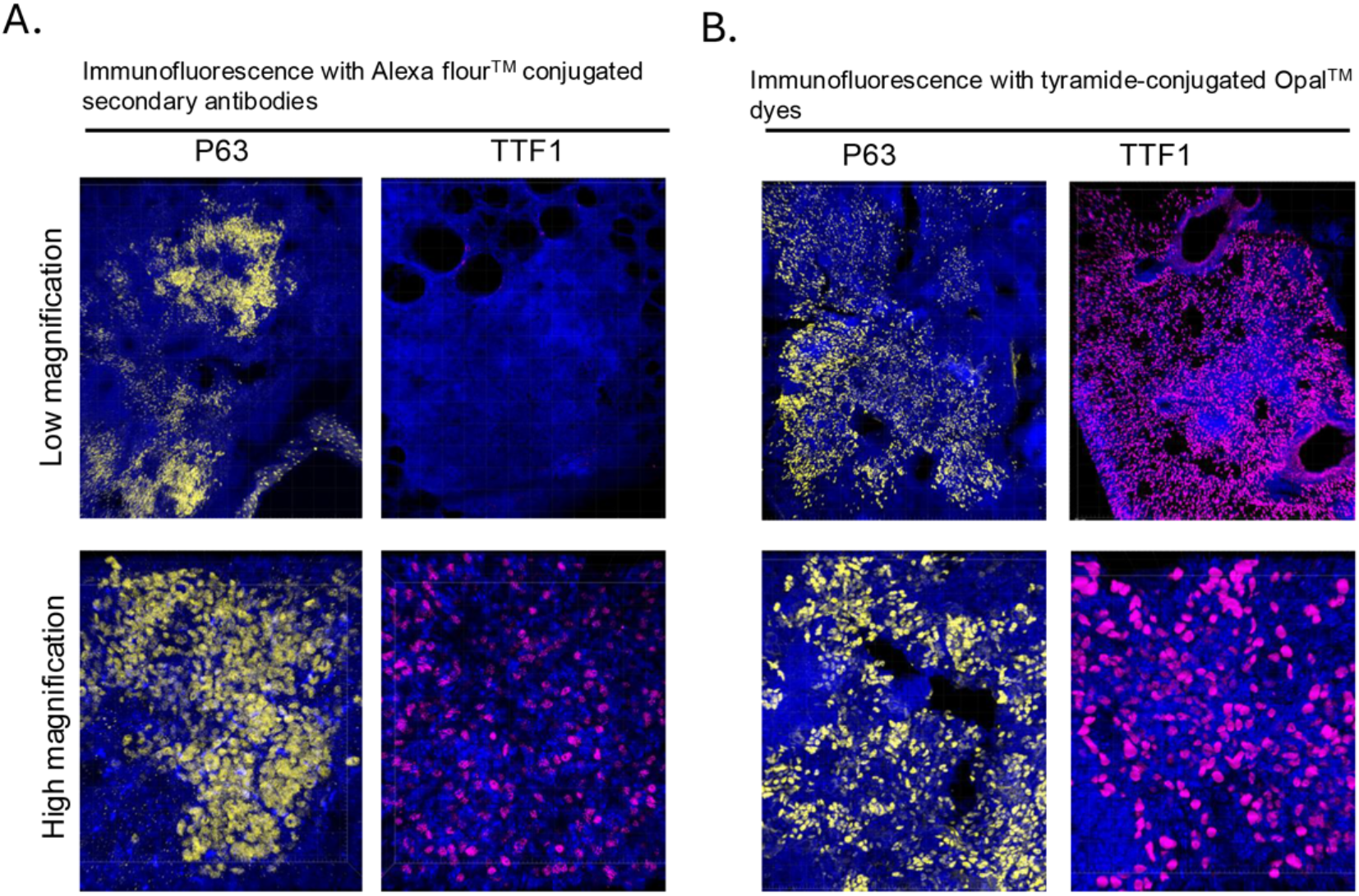
Comparison of immunostaining with Opal dyes™ and Alexa Fluor™ conjugated secondary antibodies. **A**. Thick sections were stained with P63 (right) and TTF1 (left) primary antibodies and Alexa fluor™ 647 conjugated secondary antibody. **B**. Serial sections were stained with P63 (left) and TTF1 (right), primary antibodies and developed with secondary HRP conjugated antibodies and Opal™ 540 dye. Upper images were acquired at low magnification (10X) and lower images at high magnification (63X). While P63 staining was visible at low magnification with both methods TTF1 staining was visible at low magnification with Opal™ dyes only. Laser power was set at 20% while acquiring images with Alexa fluor™ and 5% with opal dye™. This highlights the enhanced sensitivity, with retention of specificity of the tyramide signaling amplification (TSA) method over traditional indirect immunofluorescence on this tissue specimens. Immunostaining with opal dyes™ are stable and protected from bleaching since images are acquired with low laser power.

**Figure 5:**
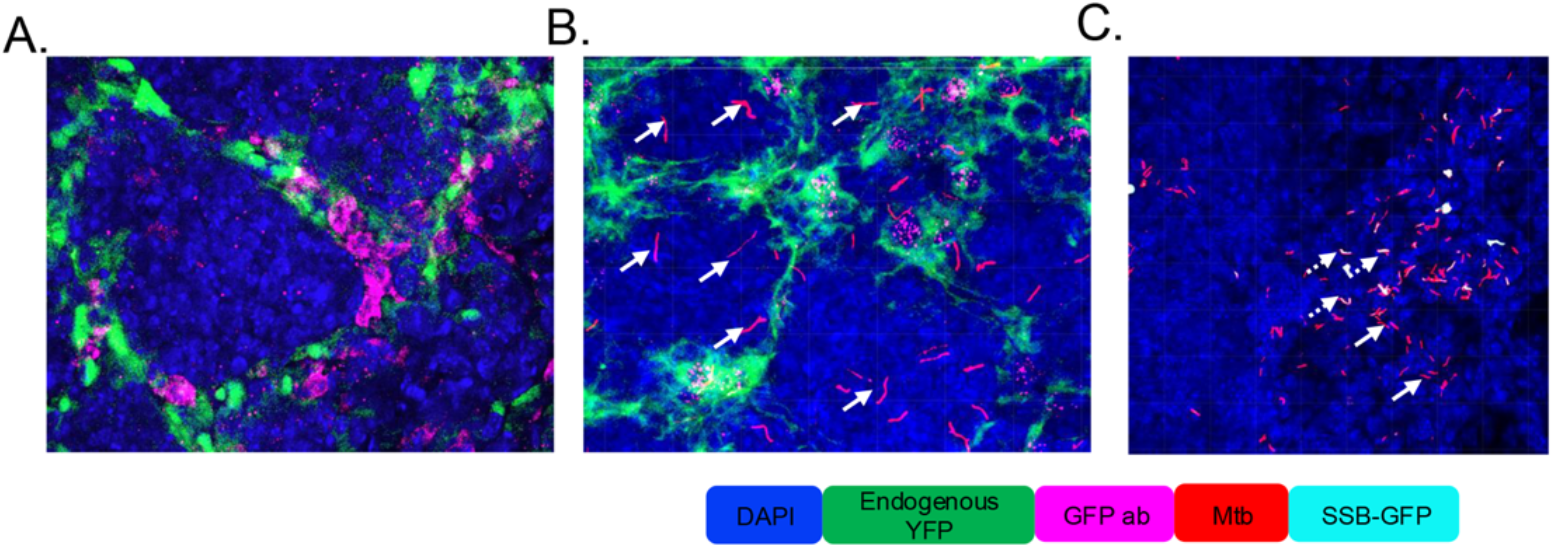
Endogenous fluorescent reporters are maintained using fluorescent multiplex immunohistochemistry (fmIHC) using Opal™ dyes. 3D confocal images of 50mm thick lung sections of reporter mice (B6.Sst1S, IFN-beta-YFP, A and B) infected with reporter Mtb (SSB:GFP, *smyc’::* mCherry). **A**. Sections were stained with anti-GFP antibody (magenta) to show the colocalization of endogenous YFP reporter with GFP antibody. **B**. Sections were stained with anti-TTF1 antibody (magenta),signal from bacterial reporter (*smyc’::* mCherry) is visible in red (white solid arrow) **C**. Endogenous YFP reporter is not seen in B6.Sst1S (non-YFP mice) signal from bacterial reporter (SSB:GFP, *smyc’::* mCherry) is visible, *smyc’::* mCherry is shown as red (white solid arrow) and replication marker SSB:GFP (Single stranded binding protein: GFP) is shown in teal; white dots represent colocalization illustrating bacterial replication inside the lung (white dotted arrow).

### Limitations

This protocol enables the use of Opal dyes ™ with thick tissue sections, resulting in enhanced signal sensitivity and reduced background when compared to indirect immunofluorescence mediated through Alexa fluor™ conjugated secondary antibodies. Since tissue sections are free floating in this protocol, there are always chances of damage to the sections while transferring samples between different incubation steps. This method is restricted to staining up to 4-5 protein markers due to the iterative 55 °C stripping buffer for 30 min, which makes the sections fragile and while transferring during washing step may result in tearing of the sections. Hence IBEX (iterative bleaching extends multiplexity) is not compatible with this method(Hickey et al., 2022). The success of this method also depends on the microscope used for image acquisition. Thick tissue sections cannot be scanned on automated imaging systems catered to thin traditional microtomy sections like the PhenoImager™. Another limitation of this method is that mouse on mouse antibodies do not work efficiently with this method (**Fig.6**).

**Figure 6:**
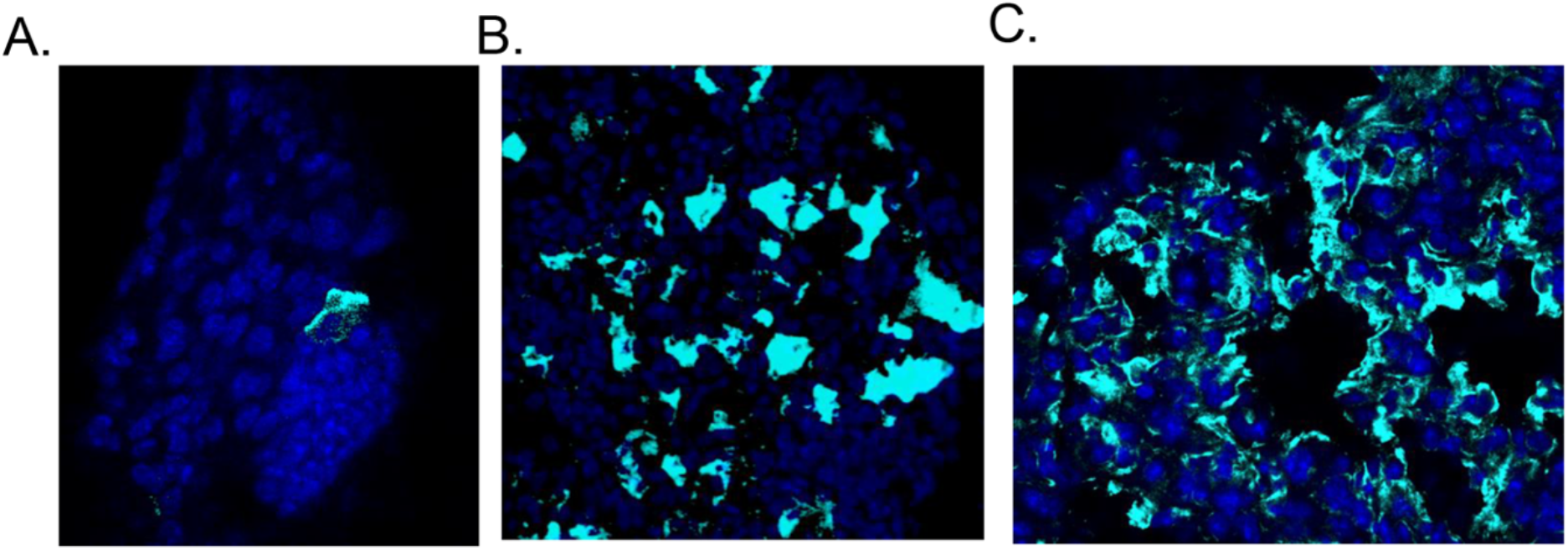
Mouse on mouse antibody doesn’t work efficiently in thick sections. **A**. Thick section was stained with mouse anti-keratin 8 antibody, incubated with rabbit IgG1 + IgG2a + IgG3 (Abcam, ab133469) at a 1:100 dilution for 30 min to block endogenous mouse IgG before adding the HRP anti-rabbit secondary antibody and developed with opal dye ™. **B**. Same mouse anti-keratin 8 antibody was used for immunostaining with FFPE (formaldehyde fixed paraffin embedded) section as discussed in (A), where it worked perfectly **C**. Thick section was stained with rabbit anti-keratin 8 antibody and developed with opal dye ™. Keratin 8 is shown in teal color. Antibody raised in rabbit work efficiently in thick section.

### Troubleshooting

#### Problem 1

##### Presence of autofluorescence

Inadequate perfusion can result in significant autofluorescence caused by red blood cells.

##### Potential solution

Perform retroorbital perfusion on live mice under anesthesia using 20-40 mL of PBS at room temperature, containing 10 IU/mL heparin. Administer the solution at a rate of 5 mL/min until the tissues are cleared of blood.

#### Problem 2

##### Tissue sections become fragile after HIER (step 4)

The integrity of tissue sections may be negatively impacted by the antigen retrieval process due to iterative exposure to high temperatures.

### Potential solution

Instead of utilizing a high temperature mediated antigen retrieval, one can permeabilize tissue sections in 1-2% of triton-X 100 for 1 h before staining. We determined this method works well serving as a sound alternative to traditional HIER (**Fig.7**)

**Figure 7:**
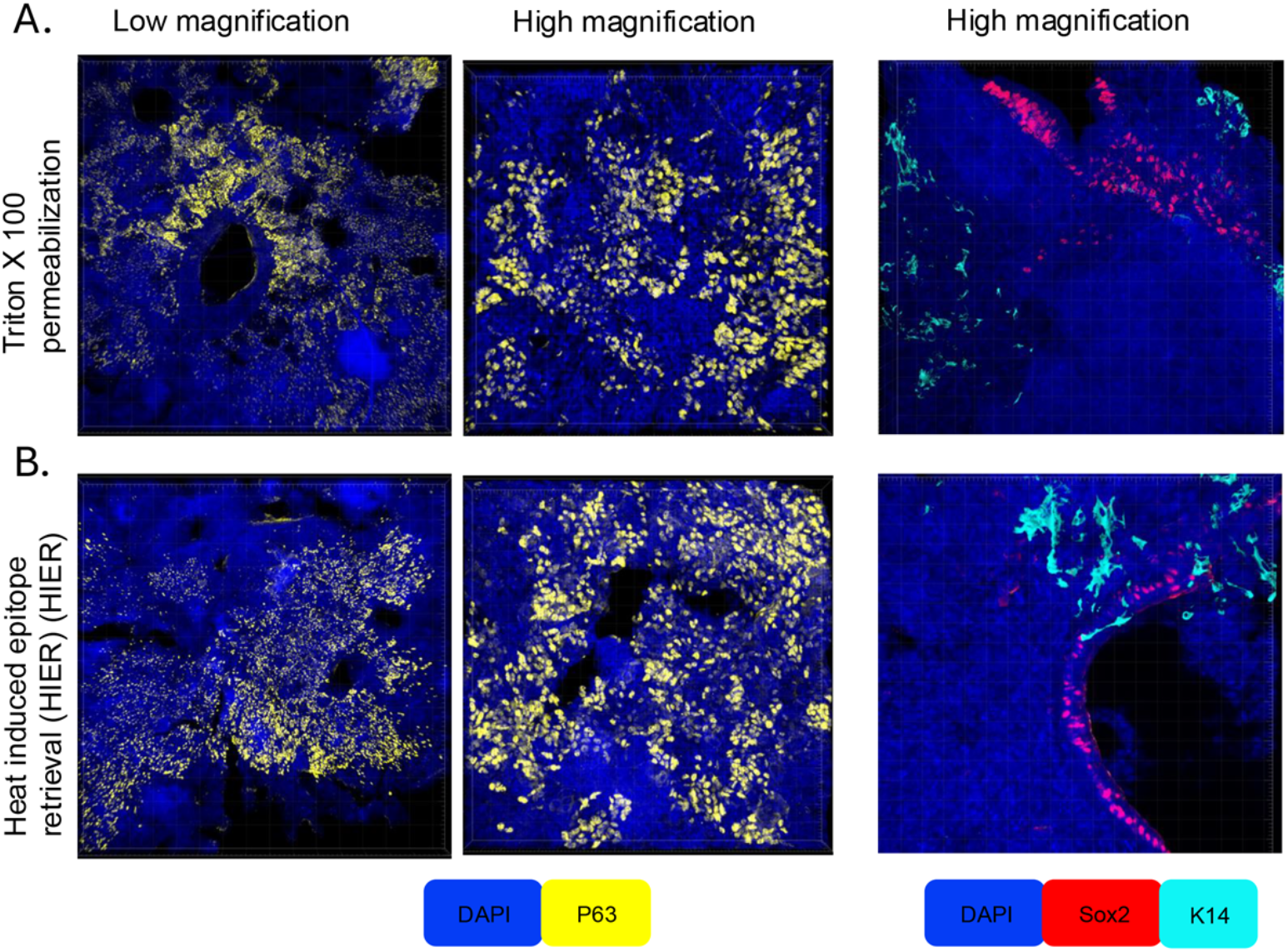
3D confocal images of P63 antibody at low and high magnification showing comparison of permeabilization and heat induced epitope retrieval (HIER) pretreatment methods. Thick sections were permeabilized by incubating in 2% triton X 100 solution in PBS at room temperature for 1h before staining process (A) or incubated in Tris/Borate/EDTA (CC1 buffer; Roche) and placed at 95°C for 30 min (HIER) before staining process (B). Both sections were stained with P63 antibody (top and middle images) utilizing TSA fluorescent immunohistochemistry using Opal dyeTM 540 (yellow) and images were acquired using a confocal microscope. Upper images are taken at low magnification (10X) and middle images at high magnification (63X). Bottom images showing 2-plex staining. Serial sections were first stained with Sox2 antibody after pretreatment, developed with Opal dyeTM 540 (red), stripped with BME/SDS stripping buffer, stained with K14 antibody and developed with Opal dyeTM 620 (teal). Both methods of pretreatment worked efficiently.

#### Problem 3

##### Markers are out of focus during imaging

Thick tissue sections need to be cleared in clearing solution before mounting to remove lipid and protein layers. We have observed clearing method significantly enhances the image quality. We have also observed issues in focusing if tissue sections are not mounted properly (refer to step 17)

### Potential solution

Clear the stained sections overnight in a clearing solution (step 16) before mounting. Dry the sections properly on a microscopic slide before adding the mounting media. Remove the excess mounting media by pressing the coverslip gently so that the distance between coverslip and tissue section should be minimum. One can also place cleared tissue sections onto a coverslip and then mount them onto a positively charged microscopic slide.

#### Problem 4

##### Bleed through between antibody targets

Bleed through between markers are seen when the pixel intensity is oversaturated bleeding into a neighboring fluorophores spectral profile, or when stripping of primary and secondary antibody complexes from a previous round is not conducted properly.

### Potential solution

Check the specificity of the primary antibody with one round of immunostaining before proceeding to a multiplex assay. An additional blocking step before addition of the primary antibody also aids in preventing non-specific binding. To confirm primary-secondary antibody complex removal, after one round of stripping, perform a control immunostaining step incubating the tissue section from the first round with a secondary antibody and Opal™ dye only. If stripping of the first round of primary-secondary antibody conjugates is successful, you should not see any signal in the control staining channel. Additionally titrate the Opal™ dye concentration as necessary before proceeding to multiplex if the pixel intensity is too strong resulting in bleed through into neighboring spectra.

#### Problem 5

##### Too strong signal

Since Opal dyes ™ undergoes tyramide signal amplification, it may result in pixel saturation and bleed through into neighboring spectra. Opal dyes™ having shorter wavelength of excitation and emission (Opal™,480, 520, & 570), are the brightest fluorophores and should be reserved for the lowest expressing proteins in your sample.

### Potential solution

Titrate the Opal dye ™ concentration in one round of staining before proceeding to multiplex.

## Notes

### Competing Interest Statement

The authors have declared no competing interest.

